# Isolation and Characterization of ΦGF1, a Morphotype C3 Bacteriophage that Infects *Escherichia coli*

**DOI:** 10.1101/627976

**Authors:** Renzo Punil, Miguel Talledo, Mayra Arcondo, Katherine Suárez, Kattya Zumaeta

## Abstract

It has been isolated a lytic bacteriophage specific to *Escherichia coli*, which can infect at least one different bacterial group. Phage ФGF1 was isolated from a wastewater treatment plant. It is resistant to the effect of chloroform and is stable at 40 and 50 °C. In addition, it is stable in the range of pH 5-8. Its host range is wide, infecting even strains from another genus such as *Shigella*. The one-step growth curve yielded a short latent period of 15 minutes and a burst size of 85 PFU per infected cell. Under the electron microscope, this phage presents the C3 morphotype, extremely rare among members of the Podoviridae family. Phage ФGF1 shows some characteristics that could be considered useful in biocontrol applications against *E. coli*. Keywords: Bacteriophage, *Escherichia coli*, morphotype C3, *Podoviridae*.

**IMPORTANCE:** Wastewater throughout the world is a heavy carrier of potential pathogens that live in their environment along with other biological agents, such as bacteriophages, which play a controlling role of the bacterial populations there, as in soil. The description of the diversity of such bacteriophages is of paramount importance since they could be used to intentionally reduce or remove those pathogens from that environment. Our work describes a bacteriophage that lives primarily in this type of water.

## INTRODUCTION

Bacteriophages are viruses that infect bacteria but no other cellular forms and whose discovery is attributed both to British microbiologist Frederick W. Twort (1915) and to French-Canadian bacteriologist Felix d’Herelle in 1917 (X. Wittebole *et* al., 2014; D.H. Duckworth, 1976). While the initial interest of the scientific community about bacteriophages was its use as tools to fight bacterial diseases, over time this idea was left aside and its study presented new perspectives, becoming the onset of new techniques today considered elemental in molecular biology, thanks to Schlesinger studies, as well as those of Hershey and Chase (R. Sharp, 2001). Furthermore, although it is very common for bacteriophages to have a high specificity for their host, there are some cases in which the host range is broad and could indicate some utility in biocontrol applications and their ability to model microbial distribution in natural environments (B. Koskella and S. Meaden, 2013; Ross *et al*, 2016). In any case, this specificity depends on the ability of the virus to bind to bacterial receptors (J. Bertozzi *et al*, 2016).

*Escherichia coli* is a gram-negative rod-shaped bacterium that can cause a wide range of diseases, mainly in the intestinal tract, although pathogenic strains have also been found that affect the urinary tract, the bloodstream and the central nervous system (J.B. Kaper *et al*. 2004; M.A. Croxen and B.B. Finlay, 2010).

Some strains of *E. coli*, in addition to produce cytotoxins, can also produce other virulent factors such as intimin and hemolysin (P.K. Fagan *et al*., 1999); for these reasons *E. coli* is considered a pathogen capable of causing diarrhea outbreaks, hemolytic uremic syndrome, hemorrhagic colitis and dysentery, mostly in children (M.A. Croxen *et al*., 2013) and many of these diseases are transmitted by food.

Foodborne diseases are a threat to public health worldwide, mainly because many of these bacteria have become more aggressive and resistant to antibiotics (O.A. Odeyemi and N.A. Sani, 2016). *E. coli* is among the most common pathogenic bacteria that are transmitted by food (T. González and R. Rojas, 2005), in addition generates great economic losses in the food industry, where antibiotics cannot be used because they generate bacterial resistance and physical and chemical treatments to inactivate these bacteria affect the organoleptic properties of food (P. Garcia *et al*., 2010). It is why non - thermal alternatives are sought for the elimination or reduction of the bacterial load, without affecting the organoleptic characteristics of the food (M. Somolinos *et al*., 2008).

The development of the use of bacteriophages for therapy is long term, since it requires many regulations in the western world and therefore many companies have opted for the application of these viruses in the field of food safety (T.K. Lu and M.S. Koeris, 2011), as the proteins derived from these viruses (endolysins) for the biocontrol of pathogenic bacteria in foods, without altering their organoleptic properties (P. García *et al*., 2010). Added to this, several studies have shown that bacteriophages can lyse multidrug-resistant bacteria taking advantage of the fact that their mechanism of action is different from that of antibiotics (I. Haq *et al*. 2012; C. Verraes *et al.*, 2013; A. Nilsson, 2014).

Bacteriophages have already been used for the reduction or elimination of the bacterial load of different pathogens in different types of food, but not all bacteriophages are efficient eliminating in the process of bacterial reduction or elimination. Therefore, it is necessary to know the microbiological, physicochemical and molecular characteristics of the bacteriophage to be used (G.A. Gonçalves et al., 2015).

The usefulness of bacteriophages depends both on their own biological properties and on the environment where they will be used, so the goal of this study was the determination of the microbiological and physicochemical properties of the lytic bacteriophage ФGF1 that infects *Escherichia coli.*

## RESULTS

### Isolation and purification of bacteriophage

Bacteriophage ФGF1 was isolated from a positive sample in broth (Figure 1A). Successive double soft layer agar assays led to the isolation of a pure phage, and through the spot test it was possible to demonstrate the lytic activity of this phage against *Escherichia coli* ATCC^®^ 25922 ™ (Figure 1B).

**Figure 1.**
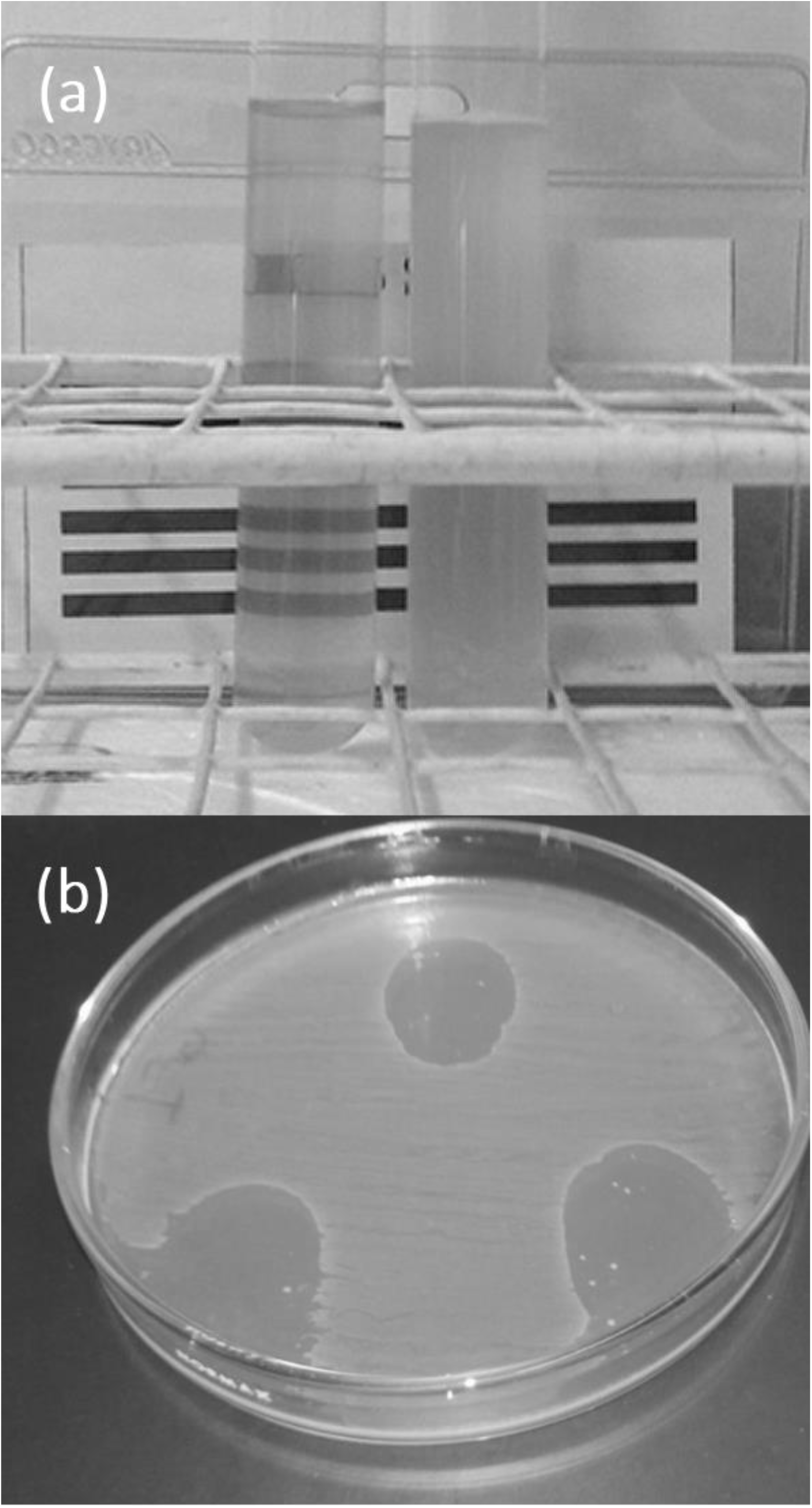
(A) Broth clearance after 8 hours incubation suggests the presence of bacteriophages. (B) Positive “spot test” of ФGF1 against *Escherichia coli* ATCC® 25922™, where three lysis spots are observed.

### Sensitivity to chloroform

The viability of phage ФGF1 was not affected after 1 hour of exposure to chloroform compared to the control test, in both cases a very similar PFU/mL value was observed (Figure 2).

**Figure 2.**
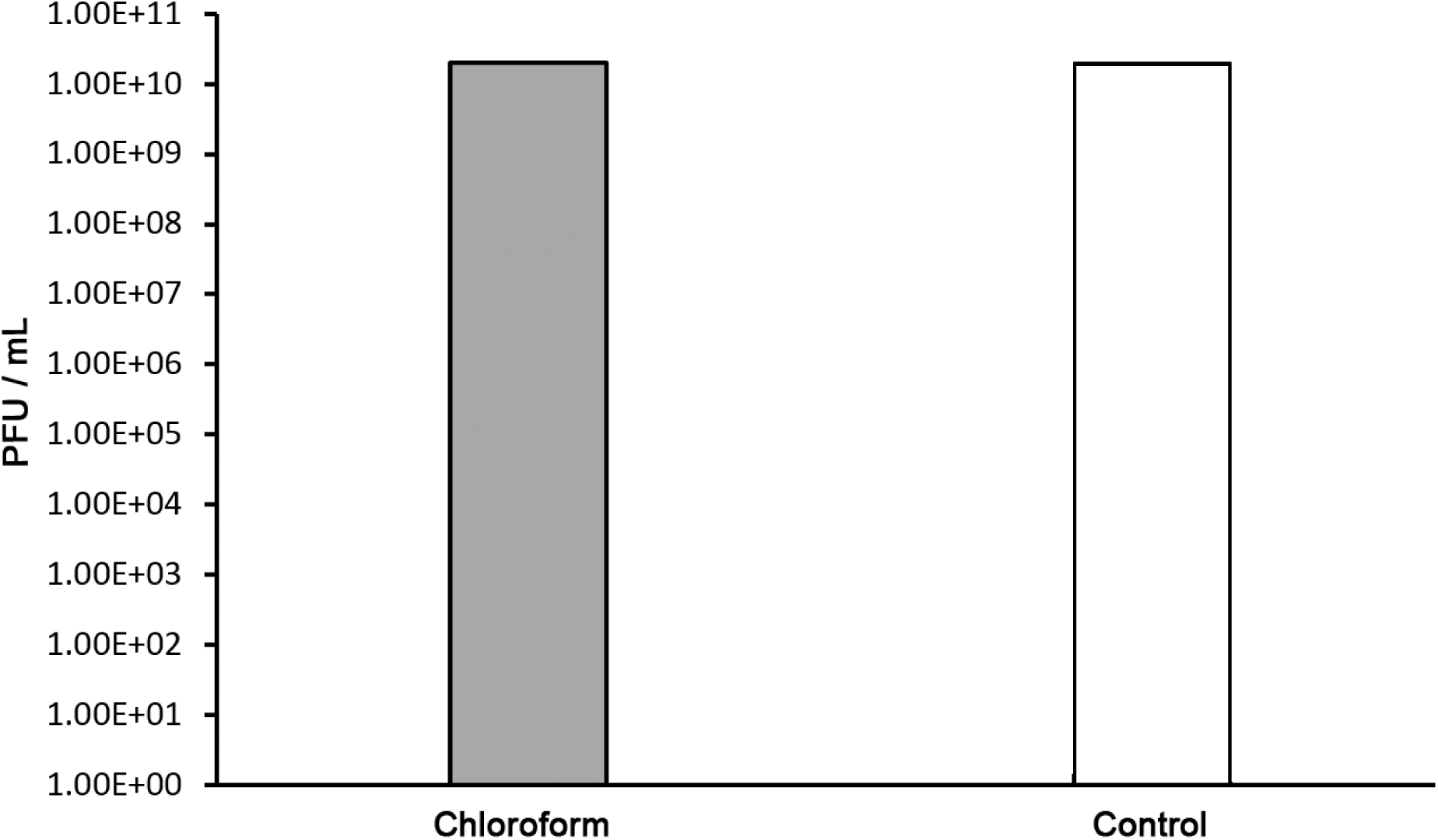
The effect of chloroform on ФGF1 stability after 60 minutes of exposure.

### Thermal stability

Thermal stability of phage ФGF1 is maintained at temperature ranging from 40 to 50 °C for up to 1 hour, and the phage is completely inactivated after 30 minutes at 70 °C and 5 minutes at 80 °C, as showed in Figure 3.

**Figure 3.**
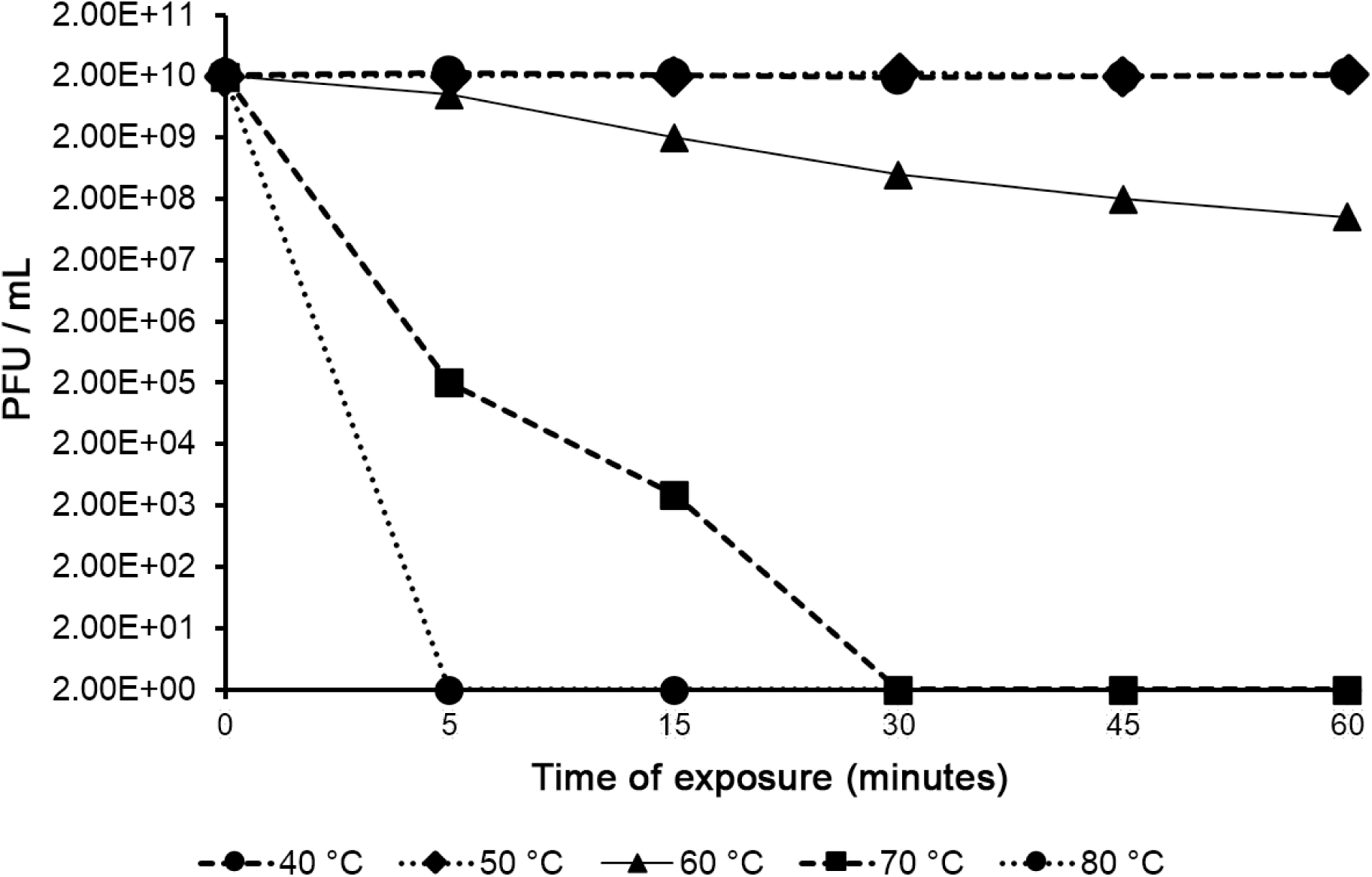
Effect of temperature on bacteriophage stability checked at 40 °C, 50 °C, 60 °C, 70 °C and 80 °C after 5, 15 30, 45 and 60 minutes.

### pH stability

ФGF1 is stable in the range of pH 5 to 8, while at pH of 9 and 10 its viability decreases up to 54 %. Acid environments of pH 3 and 4, significantly affects the viability of the phage, causing a 3-log decrease (Figure 4). However, none of the assayed pH variations completely inactivates ФGF1 after 1 hour of exposure.

**Figure 4.**
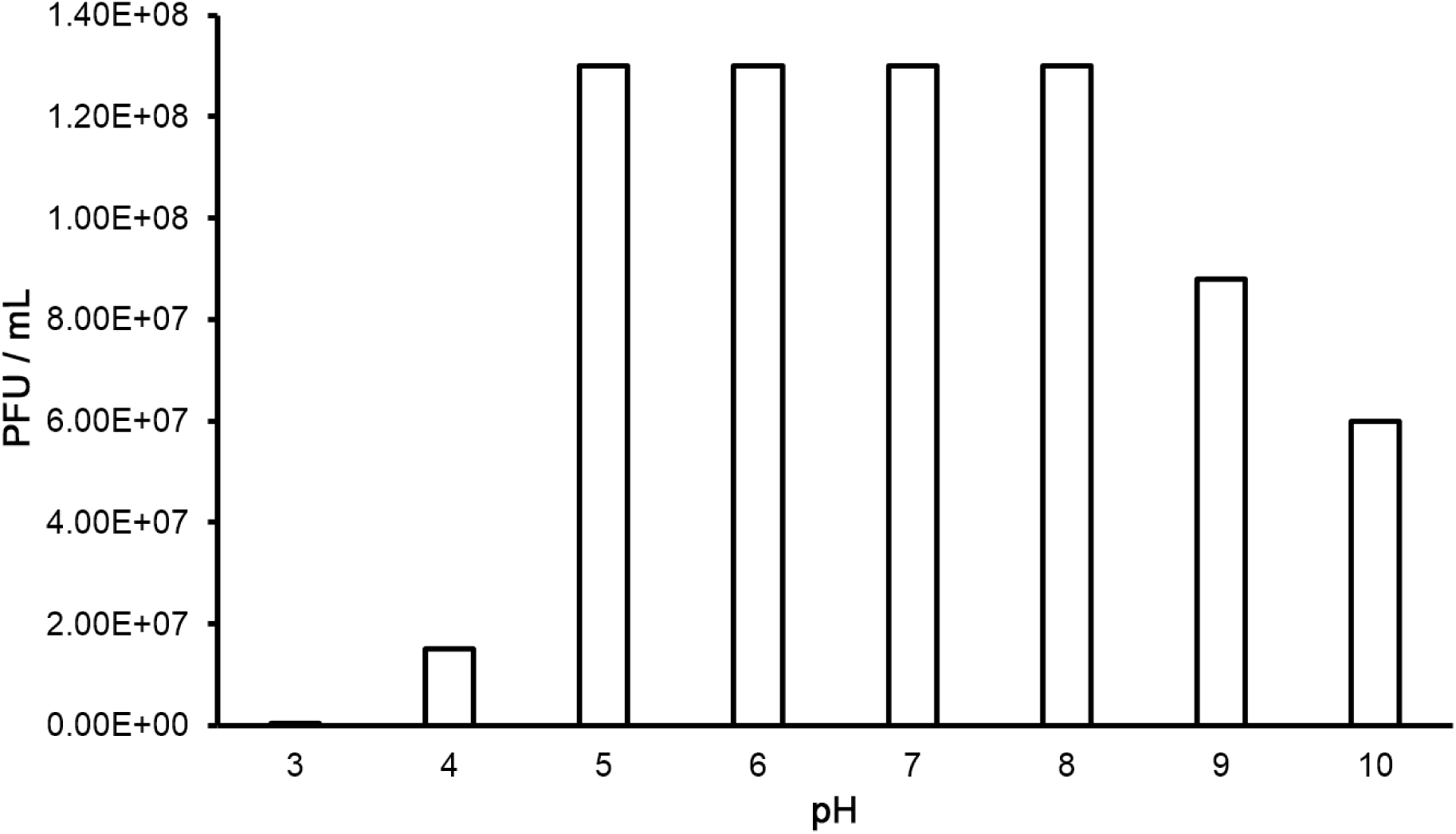
Effect of pH on bacteriophage stability. ФGF1 lysate was treated at different pH values (3, 4, 5, 6, 7, 8, 9 and 10) for one hour at 37 °C and followed by calculating phage titer by the double agar overlay technique.

### Determination of the lysis spectrum of ФGF1

Thirty one strains were evaluated, among them, other enterobacteria and gram-positive bacteria. Sensitive strains to phage ФGF1 were *E. coli* GF1, GF2 and EC3 (laboratory wild type strains), along with *Escherichia coli* ATCC^®^ 13706™, *Escherichia coli* ATCC^®^ 25922™ and *Shigella sonnei* ATCC^®^ 25931™.

### Multiplicity of infection (MOI) and one-step growth curve

Phage ФGF1 showed an optimal MOI of 0.01, a value that was used to carry on the one-step curve assay. A burst size of 85 PFU/cell was observed with a latent period of 15 minutes (Figure 5).

**Figure 5.**
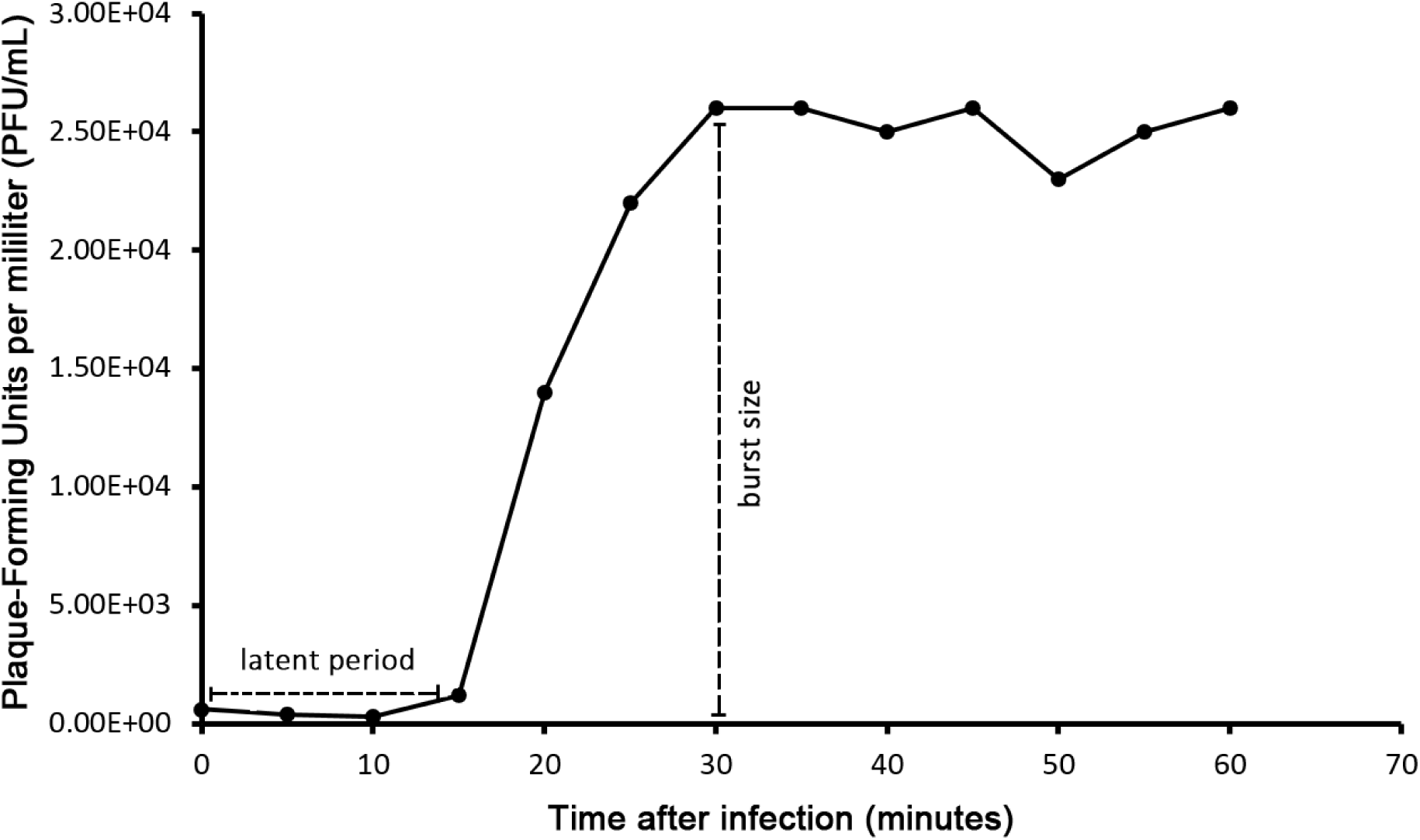
One-step growth curve of phage ФGF1. The graph shows the plaque-forming units at different times (in minutes). The length of the latent period is 15 minutes and the burst size was estimated to be 85 PFU per each infected cell.

### Transmission electron microscopy

Ultrastructure of phage ФGF1 was studied by TEM. According to this, ФGF1 belongs to the Podoviridae virus family, characterized by an elongated head and a very short tail (Figure 6). In addition, morphological similarities with the viral genus *Phieco32virus* are observed, including a 125 nm length and a 41 nm width a capsid and a tail with a length of 20 nm and a width of 11 nm.

**Figure 6.**
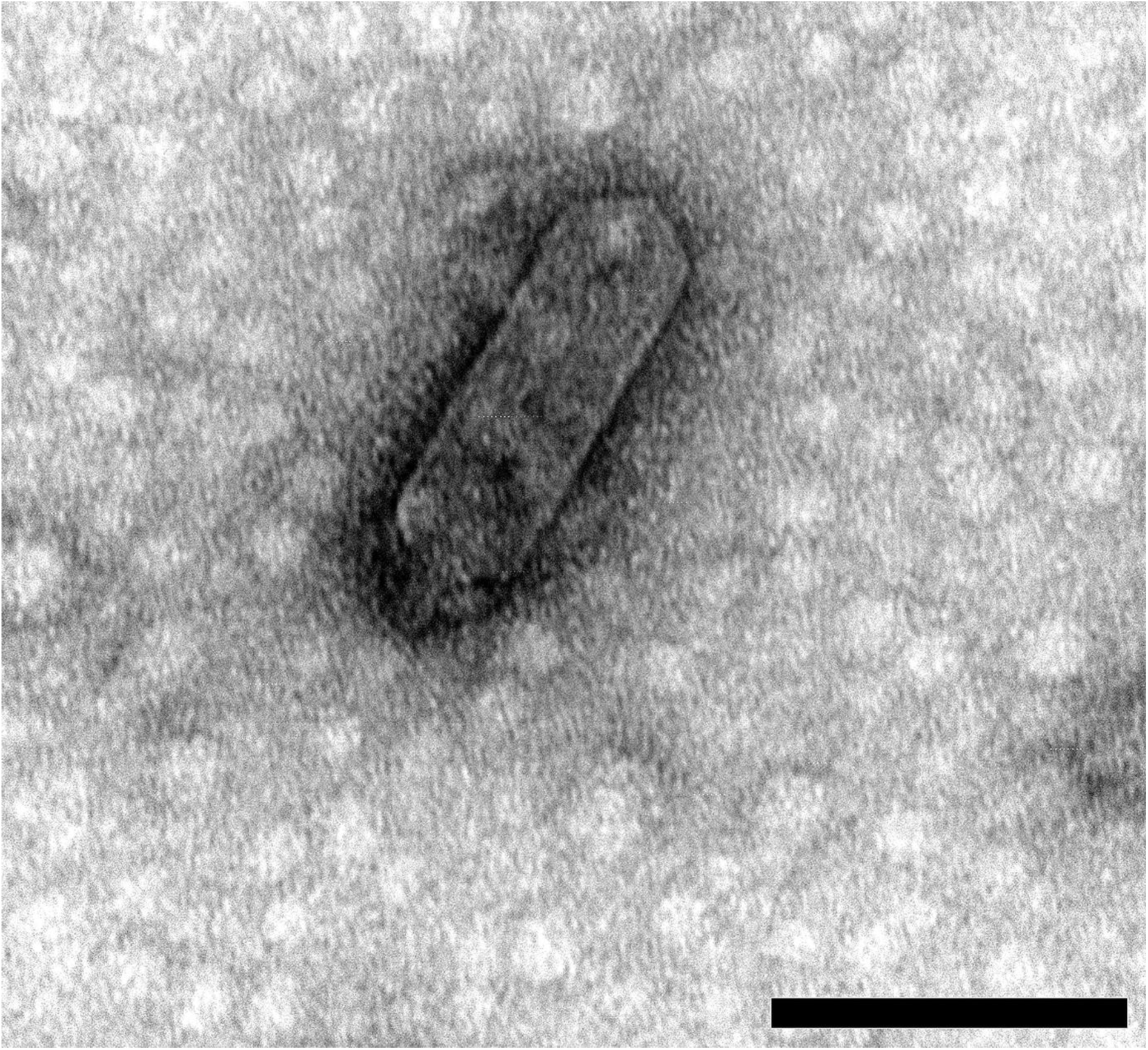
TEM image of ΦGF1. Morphology corresponds to the C3 morphotype from the viral family Podoviridae (order Caudovirales), with a rare elongated head connected to a short contractile tail by a short neck. Scale bar represents 100 nm.

## DISCUSSION

Wastewater is a good place to obtain a large number of bacterial strains, namely *Escherichia coli*, as well as non-pathogenic, pathogenic and antibiotic-resistant bacteria with great genetic diversity (E. Franz *et al*., 2015) and therefore along with them, a great diversity of specific bacteriophages. ФGF1 was isolated from the wastewater treatment plant “La Taboada”, in Lima, Peru, showing lytic activity and certain features suitable for future applications.

ФGF1 is resistant to chloroform, which is an organic compound with solvent capabilities, and although the sensitivity to chloroform is associated with the presence of lipids in the viral structure, one third of tailed phages that do not have lipids as part of the particle are sensitive to chloroform (H. W. Ackermann, 2006). In fact, chloroform can act as a destabilizing factor, and in many trials where chloroform is applied to remove the bacterial fraction after phage challenging, filtration is the preferred and only treatment (I.H. Basdew and M.D. Laing, 2014).

ФGF1 is very stable at 40 to 50 °C. This is very common in phages such as PA5oct, which infects bacteria in an optimum temperature of 37 °C (Z. Drulis-Kawa *et al*., 2014). Phage has to be adapted to its host and its environmental conditions. Va1 is a specific bacteriophage to *Vibrio alginolyticus:* the particle is stable at 20 to 30 °C, because *Vibrio* is a bacterial genus usually found in marine environments (C. Fernandez *et al*., 2017). It is pertinent to add that in certain pathogenic bacteria, such as *Burkholderia thailandensis*, temperature determines the fate of the phage inside the bacterial cell, carrying out a lytic cycle at 37 °C, but going through lysogenic cycle at 25 °C (J. Shan *et al.*, 2014).

Phage ФGF1 is stable between pH 5 and 8, as reported by N. Jamalludeen *et al*. (2007) and N.X. Hoa *et al*. (2014) who determined stability of *E. coli* specific phages in a pH range of 5 to 9. Even M.K. Taj *et* al. (2014) have found stable coliphages at pH 4. In a review by E. Jończyk *et* al. (2011), *E.* coli T7 phages can remain stable in pH ranges of 3 to 11, at very low temperatures. Very acidic pHs significantly decrease the viral concentration, but do not entirely eliminate the phage fraction.

The latent period of this phage extends to 15 minutes, close to the 19 minutes latent period reported for phage Ω8 by K. Jann *et al.* (1971), and significantly shorter in comparison to the 25 minute latent phase of phage T4 (A.H. Doermann, 1952). Although the burst size of ФGF1 was 85 plaque-forming units per infective center (PFU/IC), not as high as the 100-150 PFU/IC for T4 (A.H. Doermann, 1952), its shorter latency period and considerably high burst size make it a good candidate for future applications (M. Middelboe *et al*., 2010).

ФGF1 has a wide host range, infecting some wild strains of *E. coli, E. coli* ATCC^®^ 13706 ™ and *E. coli* ATCC^®^ 25922 ™. The particular thing about this phage is that it is not only infective for strains of the same genus, it is also infective for *Shigella sonnei* ATCC^®^ 25931™. This coincides with previous reports of common infection of *E. coli* and *Shigella* such as phage Ф24_B_ and phage CA933P (C.E. James *et al*., 2001; C. Dini, 2011). This is probably due to the fact that the genus *Shigella* is closely related to enteroinvasive *E. coli* (EIEC) (R. Lan *et al.*, 2004), as well as the direct relationship between the *E. coli* bacteriophages and the acquisition of the Shiga toxin that EIEC strains present there (A.D. O’Brien et al., 1984). Determination of the phage host range is important. P.E_1_ phage only infects some pathogenic strains of *E. coli*, which is why it is ideal for phage therapy (Z. Bibi *et al*., 2016). However, when the spectrum is broad, it can affect the intestinal natural flora, if it is used to this end (J.J. Gill and P. Hyman, 2010). On the contrary, phages with a broad spectrum, such as ФGF1, have different uses for surface decontamination and for the treatment of superficial infections (I.T. Kudva *et al*., 1999, S. O’Flaherty *et al.*, 2009) or else as food additives for preventing foodborne diseases (D. Jorquera *et al*., 2016). Currently uses include biocontrol in wastewater treatment (S.A.A. Jassim *et al*., 2016).

Morphology of ФGF1 was determined by transmission electron microscopy (TEM). It has a C3 morphotype, characterized by a capsid length that exceeds its width by several times. Phages with this morphotype are extremely rare among members of the *Podoviridae* family (Y. Li *et al*., 2012) and when they are specific for enterobacteria, they are usually related by serology and DNA homology (F. Grimont and P.A.A. Grimont, 1981). ФGF1 could belong to the genus *Phieco32virus* that has only 6 bacteriophage species, all infective for *E. coli.* This will only be confirmed by genome sequencing.

In conclusion, bacteriophage ФGF1 has a short latency period, a considerable burst size, and has a wide host range, characteristics that make it a good candidate for a diversity of biocontrol of applications *E. coli*, besides presenting an uncommon morphology to other bacteriophages already reported.

## MATERIALS AND METHODS

### Bacterial strain

The strain used as a host for the isolation of bacteriophage ФGF1 was a wild type *Escherichia coli*, isolated from “La Taboada” wastewater treatment plant, in Lima, Peru. After checking its infectivity against *Escherichia coli* ATCC^®^ 25922™, the characterization tests were carried out with this strain, which was maintained in Heart Brain Infusion broth (BHI, Merck™) at 37 °C for 24 hours.

### Phage Isolation, purification and propagation

Phage ФGF1 was isolated from wastewater prior processing at the treatment plant. A 300 mL sample was filtered through Whatman grade 1 paper. It was then filtered again in a vacuum pump (Boeco TM R-300) using 0.45 μm nitrocellulose membranes (Durapore^®^, Merck™).

To demonstrate the presence of bacteriophages in the sample and increase their number, a qualitative method was carried out; modifying what was done by S. George *et al.* (2014). Briefly, in 10 mL of BHI broth, 1 mL of the filtrate was added along with 100 μL of a log phase *Escherichia coli* ATCC^®^ 25922 ™ broth culture. A control assay was made by adding 10 mL of BHI broth, 1 mL of phosphate buffered saline (PBS) and 100 μL of the same bacterial strain. Both tubes were incubated at 37 °C for 8 hours.

The mix showing a significant clearance against the control test tube was centrifuged at 8,000 rpm for 8 minutes and the supernatant was filtered through a 0.45 μm nitrocellulose membrane. To evaluate the presence of bacteriophages, a “spot test” was applied by plating 100 μL of a *Escherichia coli* ATCC^®^ 25922 ™ overnight culture over a lawn of Tryptic Soy Agar (TSA, Merck Millipore™), and then adding 100 μL of the virus filtrate in each of three thirds of the plate.

The positive spot test filtrate was mixed with the host strain using the double agar layer technique (M.H. Adams, 1959), to produce lysis plaques with similar morphology, according to N. Jamalludeen et al. (2007). The isolated lysis plaques were transferred to phosphate buffered saline. This viral suspension was mixed with the bacterial host two more times in the same manner until the lysis plaques were uniform in shape and diameter, ensuring the purity of our bacteriophage. This last lysis plaque was cut from the agar layer, resuspended in phosphate buffered saline, filtered and mixed with a culture of *Escherichia. coli* ATCC^®^ 25922 ™ in BHI broth and incubated at 37 °C for 8 hours. Finally, this mixed solution was centrifuged at 4,400 *x g* (Thermo Scientific ST8R) for 30 minutes and filtered through nitrocellulose membranes (0.45 μm). This bacteriophage suspension was stored at −4 °C.

### Effect of chloroform

To estimate sensitivity to chloroform, C. Chenard *et al*. (2015) methodology was slightly modified. Briefly, 500 μL of phage suspension (2 × 10^10^ PFU·mL^-1^) was mixed with 500 μL of extra pure chloroform (Merck Millipore™) and kept under 250 rpm·min^-1^ for 1 hour. It was then centrifuged at 4 100 × *g* for 5 minutes and the supernatant was transferred to a microcentrifuge vial; then it was incubated for 6 hours at room temperature to remove any chloroform residue. As a control, the same procedure was performed with 500 μL of the phage suspension and 500 μL of saline solution (NaCl 0.9 % w/v). Each assay was performed in duplicate and the concentration was determined by the agar double layer technique (M.H. Adams, 1959).

### Effect of temperature

The thermal stability of the ФGF1 phage was tested at 40, 50, 60, 70 and 80 °C for 0, 5, 15, 30, 45 and 60 minutes using a phage titer of 2 × 10^10^ PFU·mL^-1^ (Z. Drulis-Kawa *et. al*., 2014). Each experiment was performed in triplicate and the phage titer was determined by the agar double layer technique.

### Effect of pH

To determine the stability of the ФGF1 phage against pH variations, pH ranging from 3 to 10 for 1 hour was assayed, slightly modifying what N.X. Hoa *et al*. (2014) proposed. 100 μL of phage suspension (1.3 × 10^8^ PFU·mL^-1^) was added to 900 μL of saline solution (NaCl 0.9 % w/v), set to a specific pH and incubated at 37 °C for 1 hour. As a control test, 100 μL of phage suspension was inoculated in 900 μL of saline solution (NaCl 0.9 % w/v) without changing pH. After incubation, each sample was adjusted to pH 7 (N.X. Hoa *et al*., 2014). Each test of pH stability was carried out in triplicate and the phage titer was determined by the double agar overlay technique (M.H. Adams, 1959).

### Host range

Bacterial susceptibility to ФGF1 phage was demonstrated by a spot test. 100 μL of an overnight culture of each bacterial strain were tested against 100 μL of phage (2 × 10^10^ PFU·mL^-1^) on TSA, spotted in three different regions of a plate, following incubation at 37 °C for 24 hours (C. Dini and P.J. de Urraza, 2010).

### Determination of multiplicity of infection (MOI) and one-step growth curve

The optimal multiplicity of infection (MOI) of the bacteriophage was determined following L. Li and Z. Zhang (2014), infecting *Escherichia* strain *coli* ATCC^®^ 25922™ at 3 different MOI (0.01, 0.1 and 1) at 37 °C for 4 hours. For the one-step growth curve experiment, 100 μL of an overnight culture of *Escherichia coli* ATCC^®^ 25922™ was inoculated to 10 mL of BHI broth and incubated at 37 °C to reach a 10^8^ CFU mL^-1^ titer (0.5 McFarland standard). Later 1 mL of this broth was mixed with 1 mL of ФGF1 phage suspension, at the optimal MOI previously determined, following incubation at 37 °C for 10 minutes and then centrifuged at 4 000 × *g* for 3 minutes. Pellet was resuspended in 2 mL of BHI broth. 100 μL of this broth were transferred to 50 mL of BHI broth and incubated at 37 °C (M. Middelboe et al., 2010). Samples (in duplicate) were taken every 5 minutes for 60 minutes and assayed by the double agar overlay technique (M.H. Adams, 1959).

### Electron microscopy of ФGF1

A concentrated phage sample was negatively stained with 2 % (w/v) uranyl acetate (pH 4.0) on a Formvar-coated copper grid and examined by Transmission Electron Microscopy (JEOL JEM-1400 Plus) (N. Jamalludeen *et al*., 2007). Phage size was determined from the average of three independent measurements.

**Table 1.** Bacterial strain susceptibility against ФGF1.

| Species | Strain | ΦGF1 |
| --- | --- | --- |
| *E. coli* GF1 | Wild type | + |
| *E. coli* GF2 | Wild type | + |
| *E. coli* AR | Wild type | - |
| *E. coli* EC1 | Wild type | - |
| *E. coli* EC2 | Wild type | - |
| *E. coli* EC3 | Wild type | + |
| *E. coli* EC5 | Wild type | - |
| *E. coli* EC6 | Wild type | - |
| *E. coli* EC7 | Wild type | - |
| *E. coli* EC8 | Wild type | - |
| *Escherichia coli* | ATCC® 13706™ | + |
| *Escherichia coli* | ATCC® 25922™ | + |
| *Shigella sonnei* | ATCC® 25931™ | + |
| *Shigella flexneri* | ATCC® 12022™ | - |
| *Salmonella* Typhimurium | ATCC® 14028™ | - |
| *Salmonella enterica* subsp. *enterica*<br>serovar *Abortusequi* | ATCC® 9842™ | - |
| *Salmonella enterica* subsp. *enterica*<br>serovar *Enteritidis* | ATCC® 13076™ | - |
| *Proteus vulgaris* | ATCC® 6380™ | - |
| *Proteus mirabilis* | ATCC® 12453™ | - |
| *Enterobacter aerogenes* | ATCC® 13048™ | - |
| *Pseudomonas aeruginosa* | ATCC® 15442™ | - |
| *Vibrio cholerae* | Wild type | - |
| *Vibrio parahaemolyticus* | ATCC® 17802™ | - |
| *Enterococcus faecalis* | ATCC® 29212™ | - |
| *Streptococcus agalactiae* | ATCC® 12386™ | - |
| *Staphylococcus epidermidis* | ATCC® 12228™ | - |
| *Bacillus cereus* | ATCC® 14579™ | - |
| *Listeria monocytogenes* | ATCC® 19114™ | - |
| *Listeria monocytogenes* | ATCC® 19115™ | - |
| *Listeria ivanovii* | ATCC® 19119™ | - |
| *Listeria innocua* | ATCC® 33090™ | - |

+ = susceptible strain
- = non-susceptible strain

## ACKNOWLEDGEMENTS

This work was supported by the Programa Nacional de Innovación para la Competitividad y Productividad (Innóvate Perú), under the Contract No. 160-PNICP-PIAP-2015, between this program and the National University Mayor of San Marcos, Lima, Peru.

At the same time, the authors thank Dr. Maurilio José Soares, main researcher from the Carlos Chagas Institute, Brazil, for his help and advice in the use of the transmission electron microscope.

